# Navigating sampling bias in discrete phylogeographic analysis: assessing the performance of an adjusted Bayes factor

**DOI:** 10.1101/2025.04.23.650183

**Authors:** Fabiana Gámbaro, Maylis Layan, Guy Baele, Bram Vrancken, Simon Dellicour

## Abstract

Bayesian phylogeographic inference is widely used in molecular epidemiological studies to reconstruct the dispersal history of pathogens. Discrete phylogeographic analysis treats geographic locations as discrete traits and infers lineage transition events among them, and is typically followed by a Bayes factor (BF) test to assess the statistical support. In the standard BF (BF_std_) test, the relative abundance of the involved trait states is not considered, which can be problematic in the case of unbalanced sampling. Existing methods to correct sampling bias in discrete phylogeographic analyses using continuous-time Markov chain (CTMC) model, often require additional epidemiological information to balance the sampling effort among locations. As such data is not necessarily available, alternative approaches that rely solely on available genomic data are needed. In this perspective, we assess the performance of a modification of the BF_std_, the adjusted Bayes factor (BF_adj_), which incorporates information on the relative abundance of samples by location when inferring support for transition events and root location inference without requiring additional data. Using a simulation framework, we assess the statistical performance of BF_std_ and BF_adj_ under varying levels of sampling bias, estimating their type I and type II error rates. Our results show that BF_adj_ complements the BF_std_ by reducing type I errors at the cost increasing type II errors for inferred transition events, while improving type I and type II errors in root location inference. Our findings provide guidelines for implementing the complementary BF_adj_ to detect and mitigate sampling bias in discrete phylogeographic inference using CTMC modelling.

## Introduction

Molecular epidemiology, the use of pathogen genomic data to address epidemiological questions, has become increasingly popular in recent years, complementing mathematical modelling, in the study of infectious disease epidemics^1^. Often, interest lies in reconstructing the dispersal history of a target pathogen to investigate the dynamic of its spread. Phylogeographic methods achieve this by combining genomic data with sampling location data to infer the ancestral locations along a phylogeny. Estimates of both the evolutionary and dispersal history are inherently uncertain. As these uncertainties can adequately be accommodated in Bayesian statistical frameworks for evolutionary analyses, several implementations for phylogeographic inference^2–4^ have become widely adopted by the field. While phylogeographic reconstructions can be conducted through a continuous (i.e., spatially explicit) diffusion^5–8^ or structured coalescent models^9–11^, today a popular method for reconstructing the dispersal history of viral lineages remains the discrete phylogeographic approach based on a continuous-time Markov chain (CTMC) model^12^. Over the last 15 years, this model has indeed been widely applied to study the spread of viruses of public health importance^13–15^. In this phylogeographic approach, sampling locations are treated as discrete characters that evolve along the phylogeny^12^. This approach was subsequently extended into a phylogeographic generalised linear model (GLM, hereafter referred to as the “CTMC-GLM” approach)^16^, allowing for explicitly evaluating the contribution of potential predictors to pathogen dispersal rates across discrete locations^17,18^.

Analogous to how the nucleotide substitution process is usually modelled, discrete phylogeographic inference using CTMC modeling^12^ employs a matrix of instantaneous rates of location exchange. Depending on whether rates from and to locations are equal or not (i.e., assuming symmetric or asymmetric transition rates), the observed sampling locations need to inform *K*\*(*K*-1)/2, or respectively *K*\*(*K*-1), parameters, with *K* being the total number of sampling locations. However, unlike nucleotide substitution models that consider multiple sites (as many as the length of the sequence alignment), each taxon in the phylogenetic tree is associated with just one location. As a result, there is an appreciable risk of model overparameterisation^12^.

To overcome this, discrete phylogeographic inference is often coupled to the Bayesian stochastic search variable selection model averaging procedure (BSSVS)^12^. This provides a Bayes factor (hereafter referred to as the standard BF annotated “BF_std_”) test to identify the subset of transition links that contribute significantly to explaining the observed distribution of locations over the tips of the phylogeny. Here, the BF_std_ support for a particular transition link is the ratio between the frequency with which that particular link is included in the model among the post-burn-in samples of the Markov chain Monte Carlo (MCMC) analysis (the posterior inclusion frequency or *a posteriori* expectation) and an *a priori* expectation (prior inclusion probability). The higher the *posterior inclusion* frequency compared to its *a priori* expectation, the higher the BF_std_ support for a non-zero transition rate between the involved locations. As it can be expected that most types of lineage transition events among locations will not occur, the default model specification assigns a 50% prior probability to the minimal rate configuration (K-1)^12^. Thus, the *a priori* expectation only depends on the number of discrete sampling locations and does not account for their relative abundances (i.e., their sampling intensity). Hence, the same prior expectation is used irrespective of the relative abundance of a location state which, in turn, may bias inferred support for the relevance of lineage transitions between some pairs of locations.

Given the importance of appropriately identifying the relevant lineage transition patterns for reliable hypothesis testing, mitigating the impact of sampling biases has been a point of attention in many studies and different approaches have been adopted. These can be summarised under four main categories: (i) conducting independent analyses based on different genomic partitions that represent different samples of the same epidemic^19^, (ii) downsampling according to local incidence estimates^20^ or to a more even number of samples by location state in the absence of such estimates^21^, (iii) incorporating data on predictors of the migration process in the CTMC-GLM approach^18,22^, and (iv) integrating individual travel history data^23^. However, almost all the above mentioned strategies rely on the availability of additional sequence or epidemiological data such as case reports, hospitalization records or flight data, which are often not readily accessible. Moreover, reaching unbiased sampling is challenging because it requires (i) a clear understanding of the extent and the intensity of the outbreak, (ii) access to the affected locations for sampling, and (iii) significant sequencing efforts^24^.

In situations where no epidemiological data is available to guide subsampling or assess how representative a dataset is of an epidemic or outbreak, Vrancken and colleagues^25,26^ proposed an alternative method to assess support for the significance of transition links that accounts for the relative abundance of samples from each location — a method that was coined the “adjusted Bayes factor” (BF_adj_)^25,26^. Concisely, instead of using the default *a priori* expectation, the authors devised an approach to incorporate information on the relative abundance of samples by location in the *a priori* expectation. For this, they rely on a discrete phylogeographic analysis during which the location states are randomly permuted, and calculate the prior odds with the inclusion frequencies from this tip-state-swap analysis.

While the BF_adj_ has been used and applied to study the spread of viruses between different locations^20,27^, its statistical performance has yet to be formally assessed. Therefore, the aim of this study is (i) to optimise the setup of the tip-state-swap analysis and (ii) to formally evaluate the statistical performance of the BF_adj_, specifically determining to what extent it can identify false positives compared to the standard approach. In other words, we seek to assess how well the BF_adj_ can identify transition events or inferred root locations for which a high BF_std_ support likely results from sampling biases. To achieve this, we use simulated epidemics of rabies virus in dogs in Morocco^28^, encompassing a total of 150 or 500 sequences representing different degrees of sampling bias. Using such a simulation framework enables to evaluate the BF_std_ and BF_adj_ type I (false positive) and type II (false negative) error rates under different conditions (i.e., different degrees of sampling bias), which allows us to here state clear recommendations regarding the use of this approach to assess the impact of sampling bias on the outcomes of discrete phylogeographic inference.

## Results

### Proportions of tip states swapped between two consecutively sampled trees

A key value to set up to conduct the tip-state-swap analysis is the relative weight of the tip-state-swap transition kernel, which dictates the expected number of permutation events during an MCMC analysis of a given length (Equation 13). The goal is to perform sufficient tip location state permutations such that the sampled trees reflect different distributions of location states. Since the tip-state-swap kernel swaps the location states of two tips at each permutation event, we studied the number of tip states swapped between two consecutives trees needed to generate a permuted “null” model while obtaining a different distribution of location states tips among the sampled trees. Therefore, we investigated the posterior indicator inclusion frequency values for all types of transition events with different numbers of expected permutations, and hence different expected proportions of tip states swapped between two consecutive sampled trees. Specifically, the latter ranged from at least 95% over at least 70%, 50%, 25%, 12.5%, 1.3%, and 0.13%, following Equation 10, for both the 150-sequence (Figure S1) and 500-sequence schemes (Figure 1A and S1). As for the standard discrete phylogeographic analysis, the MCMC length was set to 2 × 10^7^ steps for the smaller dataset and to 4 × 10^7^ steps for the larger dataset. A total of 1,000 samples were systematically collected from the converged posterior of each analysis. These analyses were done in triplicates to evaluate consistency and reproducibility.

**Figure 1.**
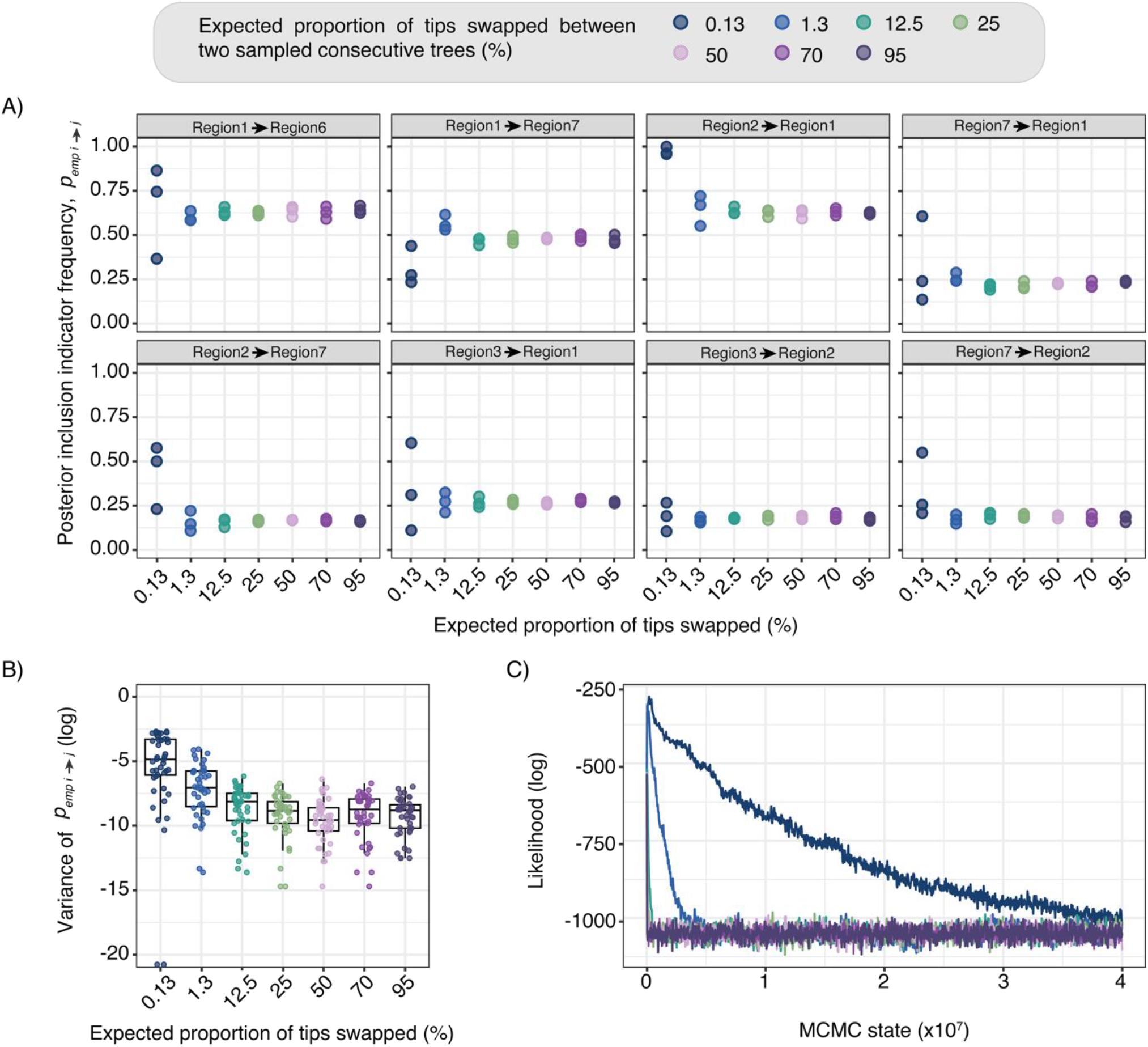
Optimising the setting of the tip-state-swap discrete phylogeographic analysis. A) Posterior inclusion indicator frequencies (*p*_*emp,i→j*_) for a set of transition events at increasing expected proportion of tip states swapped between two consecutive sampled trees. Each box represents a different transition event, from Region *i* to Region *j*. B) Distribution of the variance (log-transformed) in posterior inclusion frequencies among replicates (*n* = 3) for the different expected proportions of tip states swapped between two consecutive sampled trees. (*) For visualisation purposes only, the two values at the bottom of the plot (for the expected proportion of tips swapped equal to 0.13) were approximated to 10^−9^ instead of 0. C) Posterior traces of likelihood (log-transformed) estimates for the different runs using varying expected proportions of tip states swapped between consecutive sampled trees.

Posterior inclusion frequencies vary substantially when small proportions of tips are swapped between two consecutively sampled trees. However, a plateau is reached when having at least 25% of tip states swapped between two consecutive sampled trees, suggesting that a substantial number of permutations is necessary for achieving consistent estimates (Figure 1A and 1B). As the proportion of swapped tips increases, the variance in posterior inclusion frequencies among replicates decreases (Figure 1B). Indeed, the lowest levels of tip states swapped had significantly higher variance compared to higher levels while no significant differences were observed among proportions above 25% (Table S1). This suggests that inclusion frequencies stabilise when at least one-quarter of tip states are permuted. However, even if inclusion frequencies appear to stabilise at 25%, we chose the conservative perspective of having at least 95% of the tip swapped between two sampled consecutive trees for our downstream analysis. It is also worth noting that increasing the total number of permutations along the MCMC shortens the burn-in phase, after which the likelihood converges on that of the permuted “null” model (Figure 1C).

### Performance assessment of the BF_adj_, versus the BF_std_

Standard and tip-state-swap discrete phylogeographic inferences were run on 150-sequence and 500-sequence datasets obtained from 30 simulations conducted according to the five different sampling scenarios. To accommodate the uncertainty associated with the Bayesian phylogenetic inference but at the same time ensuring that the estimated inclusion frequencies were not impacted by potential topological differences, 100 posterior trees sampled from the standard discrete phylogeographic analysis were used as an empirical tree distribution for the corresponding tip-state-swap discrete phylogeographic inferences. Also, the latter analyses were parameterised such that at least 95% of the location states were swapped between two consecutive samples of the MCMC analysis.

We calculated the number of true positive (TP), false positive (FP), false negative (FN) and true negative (TN) transition events based on the number of transition events (i.e., number of links or branches connecting two distinct locations) along the tree averaged across the 100 subsampled posterior trees for the 150-sequence (Figure 2A and S2A) and 500-sequence datasets (Figure 2B and S2B), using a BF_std_ and BF_adj_ value of 3 as a cut-off. Our analysis showed that for both dataset sizes, the number of TP transition events identified with both the BF_std_ and BF_adj_ decreased with increasing levels of sampling bias. For the different sampling scenarios, fewer FP transition events, but also fewer TP (Table S3) and consequently higher FN transition events, were identified using the BF_adj_.

**Figure 2.**
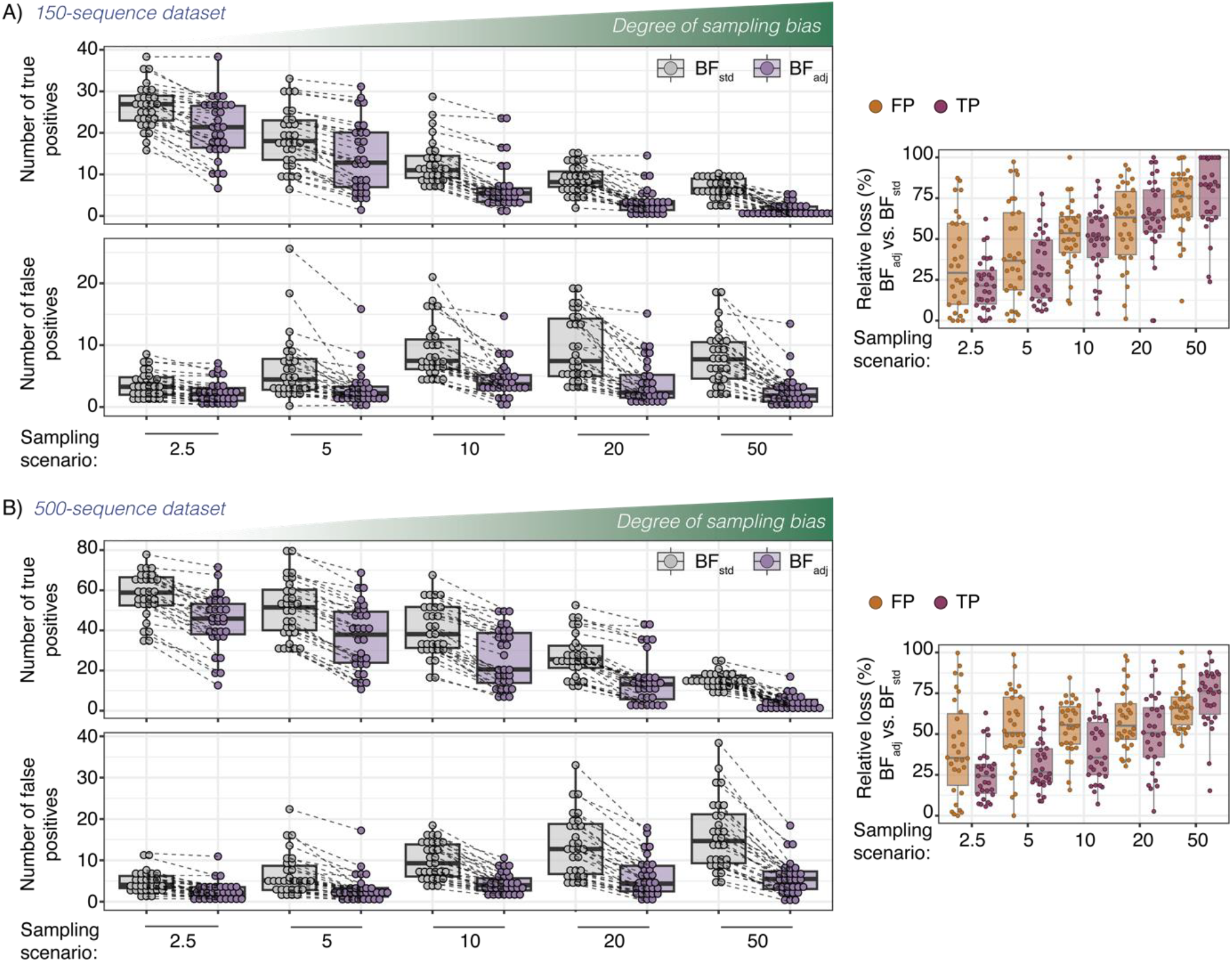
Assessment of the statistical performance of the adjusted Bayes factor (**BF**_**adj**_) in attributing statistical support to transition events inferred through a discrete phylogeographic analysis using CTMC modeling. Specifically, we here analyse the impact of the use of the BF_adj_, instead of the use of the standard BF (BF_std_) on the number of true positive (TP) and false positive (FP) transition events inferred through a discrete phylogeographic analysis under increasing levels of sampling bias. In panels A and B, we report the number of TP and FP transition events inferred by discrete phylogeographic analyses conducted on 30 datasets simulated by sampling scenario. C) Relative loss of TPs and FPs for the different sampling scenarios for the 150 (left) and 500-sequence datasets (right). For each simulation (n = 30), the relative loss of TPs (or FPs) was calculated as the percentage decrease in TPs (or FPs) when using the BF_adj_ relative to the standard BF_std_ (see the Methods section for further details).

Indeed, the relative loss of both TPs and FPs — calculated as the percentage decrease in these metrics when using BF_adj_ relative to BF_std_ — increases with the degree of sampling bias (Figure 2C). For the 150-sequence dataset, the relative loss of TPs and FPs does not differ significantly across most sampling bias scenarios (adjusted p-values > 0.05). However, in the 500-sequence dataset, the difference is statistically significant in low to intermediate bias scenarios (i.e., bias-weights of 2.5, 5, and 10), where the relative loss of FPs is greater than of TPs (with adjusted p-values ranging from < 0.0001 to 0.005). In contrast, for the high-bias scenario with a bias-weight of 20 the difference is not statistically significant (adjusted p-value = 0.178). For the most biased scenario (bias-weight of 50), the pattern inverts, with a marginally non-significant difference (adjusted p-value= 0.052), where the relative loss of TPs exceeds that of FPs (Table S3, Figure 2C).

Overall, these results suggest that BF_adj_ is more specific compared to the BF_std_, detecting fewer FPs in a comparable fashion than TPs or in some cases, more effectively than TPs, but at the cost of higher TPs loss in extremely biased datasets. While a threshold of 3 is commonly accepted as the minimum BF value for qualifying the statistical support as “positive”, higher cut-off values were also tested, and BF thresholds of 10 and 20 yielded qualitatively similar results (Figure S3).

We also assessed the performance of BF_adj_ in statistically supporting the inference of the actual root location on the tree. Across all sampling scenarios, we compared the inferred root locations obtained with those from the simulated phylogenies. Applying a cut-off value of 3, BF_adj_ consistently reduced the proportion of FPs, particularly in more biased sampling scenarios and for both dataset sizes (Figure 3). This indicates that BF_adj_ more effectively excluded incorrectly inferred root locations in a greater number of simulations compared to BF_std_. This improvement is further reflected by an increase in the proportion of TNs and a reduction in the proportion of FNs when using BF_adj_ (Figure 3). When the threshold was increased to 10 or 20, BF_adj_ led to a higher proportion of TNs while reducing FPs across most sampling scenarios for both the 150- and 500-sequence datasets (Figure 3). This effect was more pronounced for the more biased scenarios (i.e., bias-weight of 20 and 50) (Figure 3). These results suggest that applying a higher BF_adj_ threshold may improve root location inference by effectively discarding, i.e., not supporting, incorrect root location estimates.

**Figure 3.**
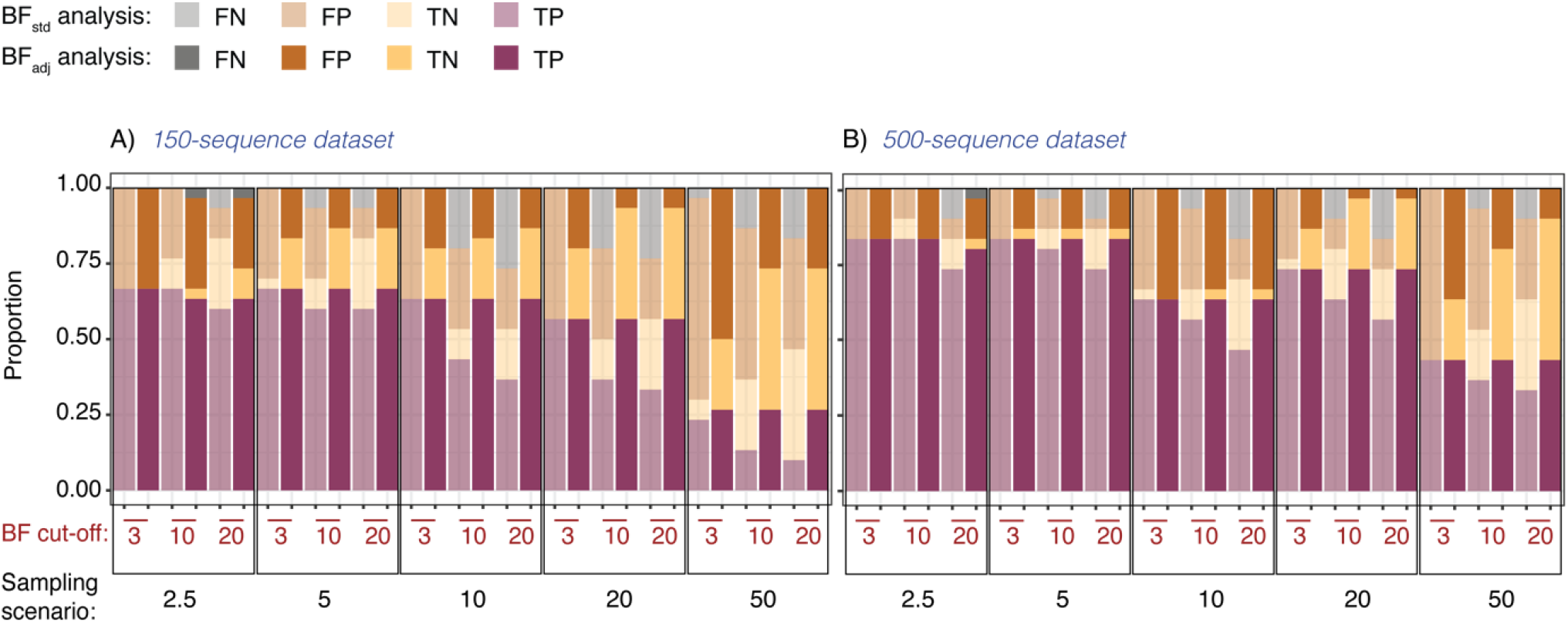
Assessment of the statistical performance of the adjusted Bayes factor (**BF**_**adj**_) in attributing statistical support to the ancestral location inferred at the root of the tree by a discrete trait analysis that uses CTMC modeling. Specifically, we here report the proportion of classification outcomes — true positives (TPs), true negatives (TNs), false positives (FP), and false negatives (TNs) — based on the ancestral location inferred at the tree root by discrete phylogeographic analyses conducted on 30 datasets simulated by sampling bias scenario, as well as for the 150- and 500-sequence datasets. We also report the results obtained when considering different cut-off values for both the BF_adj_ and BF_std_ to support an ancestral location inferred at the root node. Within the barplots, transparent and non-transparent bars correspond to results obtained using the BF_std_ and BF_adj_, respectively. The x-axes present two levels: the first level indicates the different cut-off values considered for BF_std_ and BF_adj_ (in red), and the second level refers to the different sampling scenarios.

## Discussion

In the first part of the study, we outline guidelines for setting up the tip-state-swap discrete phylogeographic analysis needed to estimate BF_adj_ support for pathogen dispersal between (discrete) locations. Analogous to the tip-date randomisation procedure used to assess the significance of the temporal signal associated with a sequence alignment^29^, the tip-state-swap analysis incorporates a permutation procedure that randomly shuffles the location states at the tips of the tree. The tip location state permutation frequency is determined by the relative importance of the tip-state-swap transition kernel. Our exploration analysis suggests that a relatively low percentage of ∼25% of tip states swapped can already achieve a proper permutation of tip states leading to robust estimates of posterior inclusion frequencies (Figure 1A and 1B). However, we recommend ensuring that at least 95% of tip states swap between consecutively sampled trees to generate a robust null permuted model where sampled trees reflect a different distribution of location states across different discrete phylogeographic analysis settings that might have not been tested here. To achieve this threshold, the weight of the tip-state-swap transition kernel must be manually adjusted, accounting for the number of taxa and the length of the MCMC, as described in Equation 13.

In the second part of the study, we use simulated RABV genomic datasets associated with different levels of sampling bias to evaluate the performance of the BF_adj_ in attributing statistical support to transition events inferred through a discrete phylogeographic analysis. Specifically, the use of simulated data enabled calculating classification outcome metrics — proportions of true positive, true negative, false positive and false negative inferred transition events among locations — to test the performance of the BF_adj_ versus the BF_std_ in the presence of varying degrees of sampling bias. We demonstrate that under all the different sampling conditions, the BF_adj_ results in less FP but also fewer TP and more FN inferred transition events. For the 150-sequence dataset, this loss of TPs was comparable to the one of FPs, meaning that with the BF_adj_, we are identifying less TPs and FPs in a similar way. For the 500-sequence dataset, the loss of FPs was greater than TPs in low to intermediate bias scenarios (i.e., bias-weights of 2.5, 5, and 10) and it gets inverted for the most biased scenario (bias-weight of 50; Table S2, Figure 2C).

Notably, for the scenarios representing a high degree of sampling bias (i.e., bias-weight of 50), the number of TPs detected with the BF_adj_ drops substantially in comparison to less biased sampling scenarios, highlighting a limitation of the BF_adj_. When the overrepresentation of certain locations in the dataset is too high, as is the case for the sampling scenario with a bias-weight of 50 (where ∼50% of the sequences come from Region 3 and ∼50% from Region 4), the BF_adj_ tends to classify transition events as non-supported. Hence, while the BF_adj_ appears to efficiently reduce FPs, it might come at the expense of being more conservative and losing sensitivity, particularly in more biased sampling scenarios. In other words, the reduction in FPs observed with BF_adj_ corresponds to a lower rate of type I errors, meaning fewer incorrect inferences of supported transition events. However, this comes at the cost of a higher number of type II errors, as the BF_adj_ also leads to more FNs (Figure S2), particularly in highly biased scenarios, indicating a reduced ability to detect TP transition events.

In the third part of the study, we used the same simulated datasets to evaluate the statistical performance of the BF_adj_ in attributing statistical support to the ancestral location inferred at the root of the tree. In this context, the use of the BF_adj_ instead of the BF_std_ results in fewer FPs although not necessarily less TPs. This suggests that while the BF_adj_ might not always support the correct root location, it is more effective at discarding incorrectly inferred root locations compared to the BF_std_. Furthermore, increasing the BF_adj_ threshold to 10 or 20 can reduce even further the risk of FP while increasing the numbers of TNs in comparison to using a BF_adj_ threshold of 3, indicating improved specificity in the inference of the root location. Hence, for the root location inference, BF_adj_ leads to a lower type I error rate and lower type II error rate, particularly for the highly biased sampling scenarios.

Inferring both the root location and transition events among discrete sampling locations are primary objectives when conducting a discrete phylogeographic analysis as they provide clues to the geographic origin of an organism and its patterns of spread. However, a significant challenge in most genomic epidemiological studies is the lack of complementary data to inform subsampling strategies, making it unclear how well the available genomic data mirrors the organism’s abundance across different locations. This limitation almost inevitably leads to unbalanced datasets that can, in turn, affect the reconstructed patterns of spread through a phylogeographic reconstruction. The adjusted Bayes factor BF_adj_, a modification of the standard Bayes factor support computed following a discrete phylogeographic analysis, attempts to incorporate information on the relative abundance of samples by location. The key notion is that when a particular transition link is strongly supported in both the standard and tip-state-swap discrete phylogeographic analysis, the associated statistical support may result merely from the high relative abundance of the involved locations. In those conditions, transition events among those locations or the location inferred at the root of the tree will be poorly supported by the BF_adj_. In our study we show that the BF_adj_ can complement the standard BF_std_ by improving (i) on the identification of transition events by reducing type I errors, but at the price of a higher number of type II errors, particularly in highly biased scenarios; and (ii) the ancestral location inferred at the root by lower type I error rate and lower type II error rate.

When epidemiological and other non-genomic data are available to inform the subsampling of the genomic dataset or provide insights into the representativeness of the sampled location in the dataset, the use of the BF_std_ should be prioritised over the BF_adj_ as it can be too conservative. For instance, using local incidence estimates to inform the (sub)sampling allows estimating transition rates among locations that can reflect the impact of spatial heterogeneity in the density of the organism’s distribution. However, when such external data are not available to inform the (sub)sampling, computing BF_adj_ supports can then represent an alternative approach allowing researchers to challenge their results in assessing whether observed patterns are due to the overrepresentation of genomes from certain locations or not, enhancing the robustness of discrete phylogeographic inference.

## Methods

### Simulated data

We used 30 simulated epidemics of rabies virus (RABV) in dogs in Morocco from the study of Layan and colleagues^28^. The epidemics were simulated across seven locations using a stochastic metapopulation model. The connectivity matrix for this model was parameterised based on human population mobility, estimated by fitting a radiation model to human population density data from the WorldPop database (https://www.worldpop.org/). Each simulated epidemic began with the introduction of a single case and culminated in at least 60,000 cases over a simulation period of 20 to 30 years^28^.

RABV genomes associated with each case were simulated along the resulting transmission chains using the Hasegawa-Kishino-Yano (HKY) substitution model^30^, taking as a starting point the complete RABV genome isolated from a dog in Morocco in 2013 (GenBank accession number KF155001.1). Viral genomes were subsequently subsampled starting one year into the epidemic to create datasets comprising either 150 or 500 genomic sequences. The simulated genomes were sampled with a sampling bias with weights increasing the sampling probability of genomes from particular regions by a factor of 2.5, 5, 10, 20, or 50. The simulated datasets corresponding to bias-weights of 2.5 and 5 represented low levels of sampling bias; those with weights of 10 and 20 intermediate levels; and the simulated datasets with a weight of 50 represented high levels of sampling bias. For each simulated epidemic, we thus obtained five distinct datasets reflecting varying degrees of sampling bias under the 150 or 500 genomic sequence schemes, resulting in a total of ten analyses per simulation. From each of the simulated transmission chains, we then extracted the “true” phylogeny linking the sampled genomes by pruning all but the sampled infections.

### Discrete phylogeographic analysis

Two types of discrete phylogeographic analyses were performed using the software package BEAST 1.10.5^2^ on each of the simulated RABV datasets: a “standard discrete phylogeographic analysis” and a “tip-state-swap discrete phylogeographic analysis” where the location states at the tips were randomly permuted during the run.

#### 1. Standard discrete phylogeographic analysis

Following the work of Layan and colleagues^28^, we specified an HKY nucleotide substitution model^30^ with a strict molecular clock and a constant population size as a tree prior. To identify the subset of transition events more informative to reconstruct the dispersal history of viral lineages, the discrete analysis was extended with a BSSVS analysis through which we obtain a BF_std_ support for all lineage transition events^12^. As described in Lemey *et al*.^12^, *in the BSSVS procedure, each element (e*.*g*., *transition link) in the instantaneous rate matrix is associated with a binary indicator variable (δ). When the rate* ***λ***between locations *i* and *j* is different from zero, the indicator variable δ_*i→j*_ takes the value of “1”, and otherwise δ_*i→j*_ equals “0”. The frequency with which *r*_*i→j*_ is included in the model yields the posterior inclusion frequency (*p*_*i→j*_). As it is expected that many transition rates are equal to zero^12^ in the current BEAST 1.10.4 setup, the prior for the indicator variables δ is a truncated Poisson prior with mean η = log(2), which assigns 50% prior probability on the minimal rate configuration with *K*-1 transition rates ***λ***. The prior probability is distributed over all possible non-zero rates. For the asymmetric model, this applies to *K*\*(*K*-1) transition rates while for the symmetric model *K*\*(*K*-1)/2. We implemented an asymmetric model, where the prior expectation for each transition rate ***λ***_*i→j*_ is defined as follows:

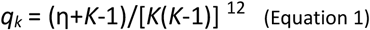

The higher the posterior inclusion frequency *p*_*i→j*_ is relative to the prior expectation *q*_*i→j*_, the more likely transition events between locations *i* and *j* help explain the diffusion process. This appreciation is expressed by the Bayes factor, which is obtained by taking the ratio between (i) the posterior odds that the transition rate is non-zero and (ii) its prior odds:

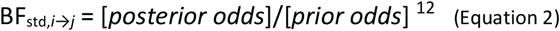

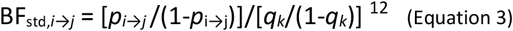

When *p*_*i→j*_ was equal to one, we avoided dividing by zero by approximating *p*_*i→j*_ as follows:

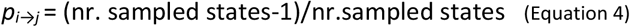

This leads to a finite value for BF_std_ which represents the lower bound for the actual support.

We also estimated the BF_std_ support for the root location. For each root location *i*, we considered the location probability *p*_*i*_ and assigned equal prior probability to all *K* locations. Therefore, the BF_std_ support for the root location *i* is given by:

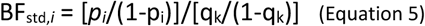

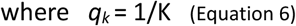

Similar as before, when the location probability for a given location *i* was equal to “1”, the location probability *p*_*i*_ was approximated following Equation 4. As BF_std_ support values represent a range of standards of evidence, thresholds of 3, 10 and 20 were considered^31^.

Analyses were run for 20 million or 40 million Markov chain Monte Carlo (MCMC) iterations, with samples collected every 20,000 or 40,000 iterations, for the 150 and 500 sequences dataset respectively. We used the BEAGLE v4.0.0 library to improve computational performance^32^. Convergence and mixing were assessed using the program Tracer 1.7.2^33^, ensuring all parameters were associated with an effective sample size (ESS) value greater than 200. After discarding the initial 10% of sampled posterior trees as burn-in, the maximum clade credibility (MCC) tree was retrieved and annotated using the program TreeAnnotator 1.10^34^.

#### 2. Tip-state-swap discrete phylogeographic analysis

For the tip-state-swap discrete phylogeographic analysis, 100 evenly sampled post-burnin trees from the corresponding standard discrete phylogeographic analysis were used as an empirical tree distribution^16^ to ensure that the estimated inclusion frequencies were not impacted by potential topological differences. The analyses were run for the same number of steps as the standard discrete phylogeographic analysis, and the same diffusion model was specified except that tip state locations were randomised during the MCMC simulation by including the tip-state-swap transition kernel (see the Data Availability for details on template files). To test the effect of different permutation intensities, we adjusted the transition kernel weight to achieve varying expected proportions of swapped tips between two consecutive sampled trees (e.g., 25%, 50%, 75%, and 95%).

At each permutation event, the location states of two tips (i.e., sampled sequences) were swapped. For a dataset with *n* tips, the probability *p*_*1*_ that a tip is not randomly swapped with another one is given by:

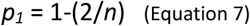

The probability *p*_*2*_ that a given tip is swapped at least once after (*np*) number of permutations is:

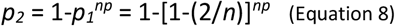

Thus, the expected number of tips *E* being swapped at least once after *np* permutations can be calculated as follows:

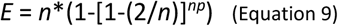

To determine the number of permutations *np* between two consecutive sampled trees needed to achieve a specified minimum proportion (*x*) of tip states swapped we do:

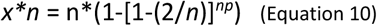

Rearranging the equation, *np* is given by:

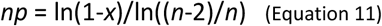

The total number of expected permutation events along the MCMC chain *tnp* is then given by

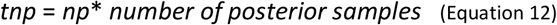

In practice, the total number of permutation events is determined by the interplay between the MCMC length and the expected frequency with which tip-state-swaps occur. The latter is governed by the relative importance of the tip-state-swap transition kernel. In this way, we can calculate the weight to assign to the tip-state-swap transition kernel for a given MCMC length and sampling frequency in order to expect at least a proportion *x* of tip states swapped between two sampled consecutive trees:

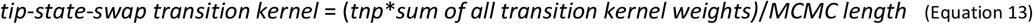

### Estimation of the adjusted Bayes Factor (*BF*_*adj*_)

Following Vrancken and colleagues^25^ as well as Chaillon and colleagues^26^, the BF_adj_ support for a transition link between locations *i* and *j* was calculated by replacing the default prior expectation based on the minimal rate configuration, which only depends on the number of sampled locations, with an empirical prior expectation *p*_*emp,i→j*_ that accounts for the relative abundance of sampled locations (i.e., sampling intensity). The *p*_*emp,i→j*_ is the mean posterior inclusion frequency for the transition link *i* to *j* obtained from the tip-state-swap discrete phylogeographic analysis. The BF_adj,*i→j*_ is then calculated as follows:

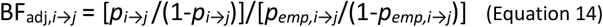

We also estimated the BF_adj_ support for the root location. Similar as for the support of the transition links, the prior expectation assigning equal *priori* probability to all *K* locations was replaced with an empirical prior expectation, which is the location probability, *p*_*emp,i*_, obtained from the tip-state-swap discrete phylogeographic analysis. Therefore, the BF_adj_ support for the root location *i* is given by:

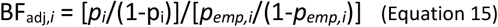

For the BF_adj_ as well, thresholds of 3, 10 and 20 were considered.

### Statistical performance assessment

To assess the statistical performances of the BF_adj_ in comparison to the BF_std_, we contrasted discrete phylogeographic reconstructions where only relevant transition links supported by the BF_std_ or BF_adj_ were considered against the “true” phylogenies derived from the simulations. Specifically, the first objective was to determine how effectively the BF_adj_ could identify transition events where strong BF_std_ support results from sampling imbalances. For this, we calculated the total number of classification outcomes, that is to say true positives (TPs), true negatives (TNs), false positives (FPs) and false negatives (FNs) across the different sampling scenarios.

For all *K*\*(*K*-1) possible transition events between the discrete sampling locations, we estimated the number of transition events in the phylogeographic reconstruction by averaging the values across 100 sampled trees. Each of these “inferred counts” were compared with the observed number of transition events between the same locations in the “true” trees (the “simulated count”). We defined the classification outcomes as follows:

1. If the BF_std_ and/or BF_adj_ ≥ threshold:
  a. if the inferred count exceeded the simulated count, the excess number were considered FPs, while the remainder were imputed to be TPs;
  b. if the inferred count matched the simulated count, the entire count was considered TPs;
  c. if the inferred count was lower than the simulated count, the inferred count was considered as the number of TPs, while the difference was scored as the number of FNs.
2. If the BF_std_ and/or BF_adj_ < threshold:
  a. if the inferred number of transition events exceeded the simulated number, the simulated number was considered as FNs and the rest as TNs;
  b. if the inferred count matched or was lower than the simulated count, the simulated count was considered as FNs.

To further evaluate the performance of the BF_adj_ under the different degrees of sampling bias, we computed the relative TPs and FPs loss for each given sampling scenario. The relative loss was calculated as the decrease in percentage of TPs (FPs) using the BF_adj_ compared to BF_std_. Specifically, for each sampling scenario and each simulation, TPs and TPs_adj_ represent the number of true positives identified with the standard and adjusted BF, respectively. The relative loss of TPs is given by:

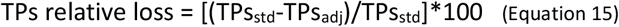

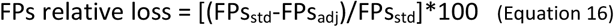

A second objective was to evaluate to which extent the BF_adj_ can help identify when the location inferred at the root of the tree was impacted by sampling bias. To this end, we looked at the location inferred at the root of the tree and its associated BF_std_ and BF_adj_ support. In practice, we again estimated the total number of classification outcomes, true positives (TPs), true negatives (TNs), false positives (FPs) and false negative (FNs) for each epidemic simulation across the different sampling scenarios by comparing the inferred root location with the location at the root of the simulated phylogenies:

1. If the inferred root location matched the simulated root location:
  a. BF_std_ and/or BF_adj_ ≥ threshold, it was counted as a TP;
  b. BF_std_ and/or BF_adj_ ≤ threshold, it was counted as a FN.
2. If the inferred root location did not match the simulated root location:
  a. BF_std_ and/or BF_adj_ ≥ threshold, it was counted as a FP;
  b. BF_std_ and/or BF_adj_ ≤ threshold, it was counted as a TN.

## Statistical analyses

All statistical analyses were performed in R version 4.2.1^35^ from the CRAN repository. To compare the variance of inclusion frequencies across different permutation levels, pairwise Levene’s tests were performed using the function “leveneTest” of the R package “car” ^36^. Resulting p-values were adjusted for multiple comparisons using “Benjamin-Hochberg” with the function “p.adjusted” from R package “stats”. The statistical significance of TPs and FPs values obtained with the BF_std_ and BF_adj_ approaches was evaluated using either paired two-tailed *t*-tests (for normally distributed data, i.e., Shapiro-Wilk test p-value > 0.05) or the paired two-tailed Wilcoxon signed-rank test (a non-parametric alternative to the paired *t*-test). The normality of distributions was assessed using a Shapiro-Wilk test as implemented in the function “shapiro.test” of the R package “stats” ^35^. As above, p-values were adjusted for multiple comparisons with “Benjamin-Hochberg” using the function “p.adjusted” from R package “stats”. Significance was determined at a p-value threshold of 0.05.

## Data Availability

R scripts and example files are available at https://github.com/FabiGambaro/BFadj. Additionally, a step-by-step tutorial on the use of the adjusted Bayes factor (BF_adj_) has been available on the BEAST X community website: https://beast.community/adjusted_bayes_factor.

## Financial Statement

FG, BV and SD acknowledge support from the *Fonds National de la Recherche Scientifique* (F.R.S.-FNRS, Belgium; including grant n°F.4515.22). GB acknowledges support from the Research Foundation - Flanders (*Fonds voor Wetenschappelijk Onderzoek — Vlaanderen*, FWO, Belgium; grant n°G0E1420N), and from the DURABLE EU4Health project 02/2023-01/2027 which is co-funded by the European Union (call EU4H-2021-PJ4) under Grant Agreement No. 101102733. SD and GB acknowledge support from the Research Foundation — Flanders (*Fonds voor Wetenschappelijk Onderzoek — Vlaanderen*, FWO, Belgium; grant n°G098321N) and from the European Union Horizon 2023 RIA project LEAPS (grant agreement no. 101094685). SD also acknowledges support from the University of Brussels (ULB, Belgium) internal fund, the BE-PIN project (TD/231/BE-PIN) funded by the Belgian Science Policy Office (BELSPO, Belgium), the ImmunReach project funded by the *Institut d’Encouragement de la Recherche Scientifique et de l’Innovation de Bruxelles* (Innoviris, Belgium), the Doctoral network VIVACE funded by the Marie Skłodowska-Curie Actions (MSCA) of the European Commission (grant agreement n°101167768), and from the European Union Horizon 2020 project MOOD (grant agreement n°874850). Computational resources have in part been provided by the Consortium des Équipements de Calcul Intensif (CÉCI), funded by the *Fonds de la Recherche Scientifique de Belgique* (F.R.S.-FNRS) under grant n°2.5020.11 and by the Walloon Region.

---

**Table S1.** Statistical comparison of the inclusion frequencies variance among different permutation levels. Significance was tested using Levene’s test on each pair, with p-value adjusted for multiple comparison using “Benjamini-Hochberg” (see the Methods section for further detail).

| Permutation level |  | 150-sequence dataset | 500-sequence dataset |
| --- | --- | --- | --- |
| group1 | group2 | Adjusted p-value | Adjusted p-value |
| 0,13 | 1,3 | < 0.001 | < 0.001 |
| 0,13 | 12,5 | < 0.001 | < 0.001 |
| 0,13 | 25 | < 0.001 | < 0.001 |
| 0,13 | 50 | < 0.001 | < 0.001 |
| 0,13 | 70 | < 0.001 | < 0.001 |
| 0,13 | 95 | < 0.001 | < 0.001 |
| 1,3 | 12,5 | 0,0469 | < 0.001 |
| 1,3 | 25 | < 0.001 | < 0.001 |
| 1,3 | 50 | < 0.001 | < 0.001 |
| 1,3 | 70 | < 0.001 | < 0.001 |
| 1,3 | 95 | < 0.001 | < 0.001 |
| 12,5 | 25 | < 0.001 | 0.039 |
| 12,5 | 50 | < 0.001 | 0.047 |
| 12,5 | 70 | < 0.001 | 0.058 |
| 12,5 | 95 | < 0.001 | 0.008 |
| 25 | 50 | 0.4065 | 0.865 |
| 25 | 70 | 0.5789 | 0.719 |
| 25 | 95 | 0.5789 | 0.660 |
| 50 | 70 | 0.7333 | 0.682 |
| 50 | 95 | 0.2518 | 0.790 |
| 70 | 95 | 0.3536 | 0.299 |

**Table S2.** Testing the differences between the number of true positives (TPs) and false positives (FPs) identified with the **BF**_**adj**_ and **BF**_**std**_. Significance was tested either with paired two-tailed *t*-tests or paired two-tailed Wilcoxon signed-rank tests with p-value adjusted for multiple comparison using “Benjamini-Hochberg” (see the Methods section for further details).

| Sampling scenario | metric | 150-sequence dataset |  | 500-sequence dataset |  |
| --- | --- | --- | --- | --- | --- |
|  |  | test | p-value | test | p-value |
| 2.5 | TPs | paired *t*-test | < 0.001 | paired *t*-test | < 0.001 |
|  | FPs | paired Wilcoxon test | < 0.001 | paired Wilcoxon test | < 0.001 |
| 5 | TPs | paired *t*-test | < 0.001 | paired *t*-test | < 0.001 |
|  | FPs | paired Wilcoxon test | < 0.001 | paired Wilcoxon test | < 0.001 |
| 10 | TPs | paired Wilcoxon test | < 0.001 | paired Wilcoxon test | < 0.001 |
|  | FPs | paired Wilcoxon test | < 0.001 | paired Wilcoxon test | < 0.001 |
| 20 | TPs | paired Wilcoxon test | < 0.001 | paired Wilcoxon test | < 0.001 |
|  | FPs | paired Wilcoxon test | < 0.001 | paired Wilcoxon test | < 0.001 |
| 50 | TPs | paired Wilcoxon test | < 0.001 | paired Wilcoxon test | < 0.001 |
|  | FPs | paired Wilcoxon test | < 0.001 | paired Wilcoxon test | < 0.001 |

**Table S3.** Testing the difference between the relative loss of FPs and the relative loss TPs associated with the use of the **BF**_**adj**_ **when conducting discrete phylogeographic analyses on 30 simulations while considering different sampling scenarios and two different dataset sizes (Figure 1A and 1B)**. Significance was tested either with paired two-tailed *t*-tests or paired two-tailed Wilcoxon signed-rank tests (see the Methods section for further detail).

| Sampling scenario | 150-sequence dataset |  | 500-sequence dataset |  |
| --- | --- | --- | --- | --- |
|  | p-value | test | p-value | test |
| 2.5 | 0.092 | paired Wilcoxon test | 0.005 | paired *t*-test |
| 5 | 0.112 | paired *t*-test | < 0.001 | paired *t*-test |
| 10 | 0.546 | paired *t*-test | 0.005 | paired *t*-tests |
| 20 | 0.522 | paired Wilcoxon test | 0.179 | paired *t*-tests |
| 50 | 0.546 | paired Wilcoxon test | 0.053 | paired Wilcoxon test |

**Figure S1.**
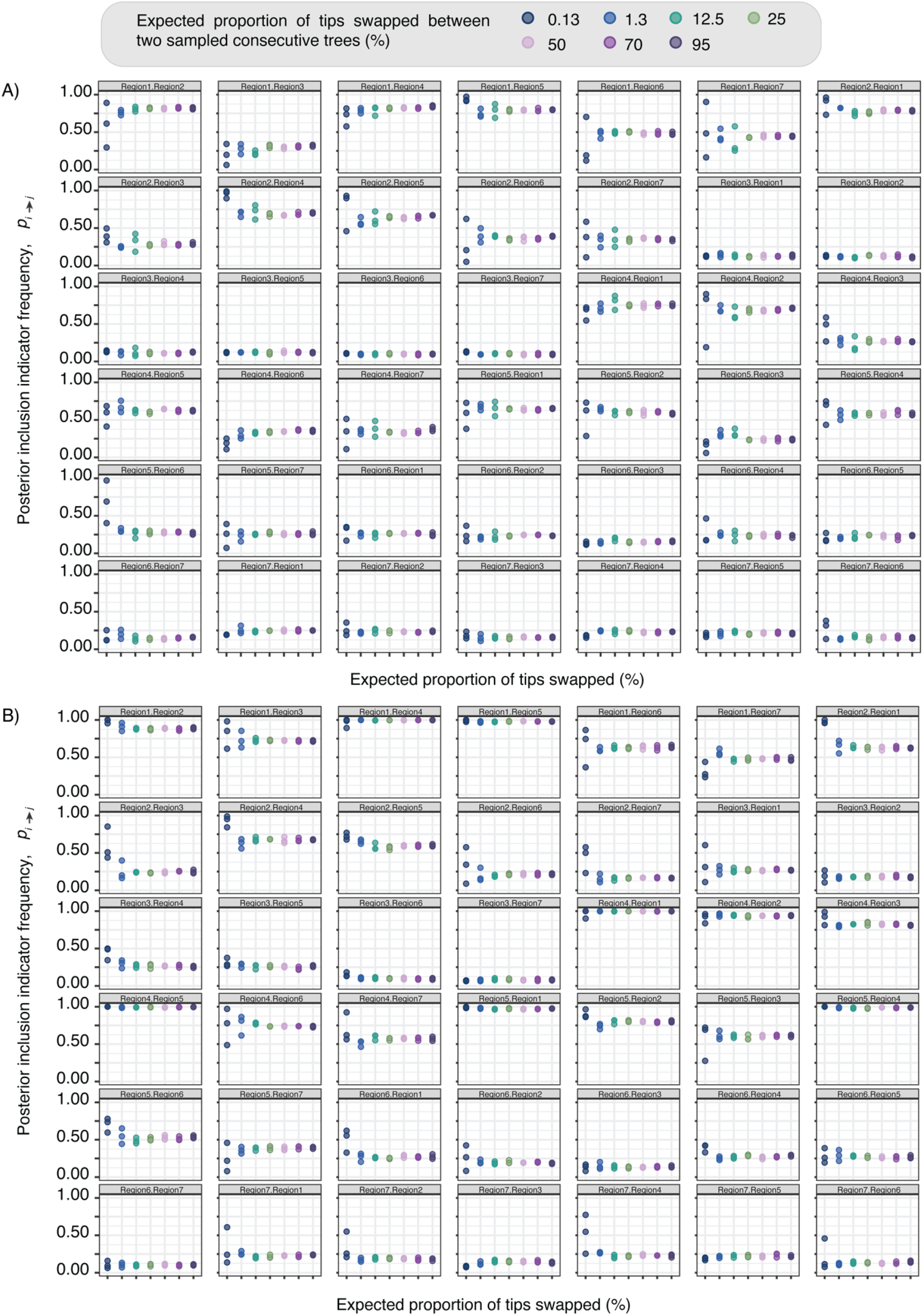
Posterior inclusion indicator frequencies (***p***_***i→j***_) for all transition events at increasing expected proportion of tip states swapped between two consecutive sampled trees. Each box represents a different transition event, from Region *i* to Region *j*. Panel A and B shows the results for the 150-sequence and 500-sequence datasets, respectively. Results are presented for three MCMC chains performed on the same simulated phylogeny.

**Figure S2.**
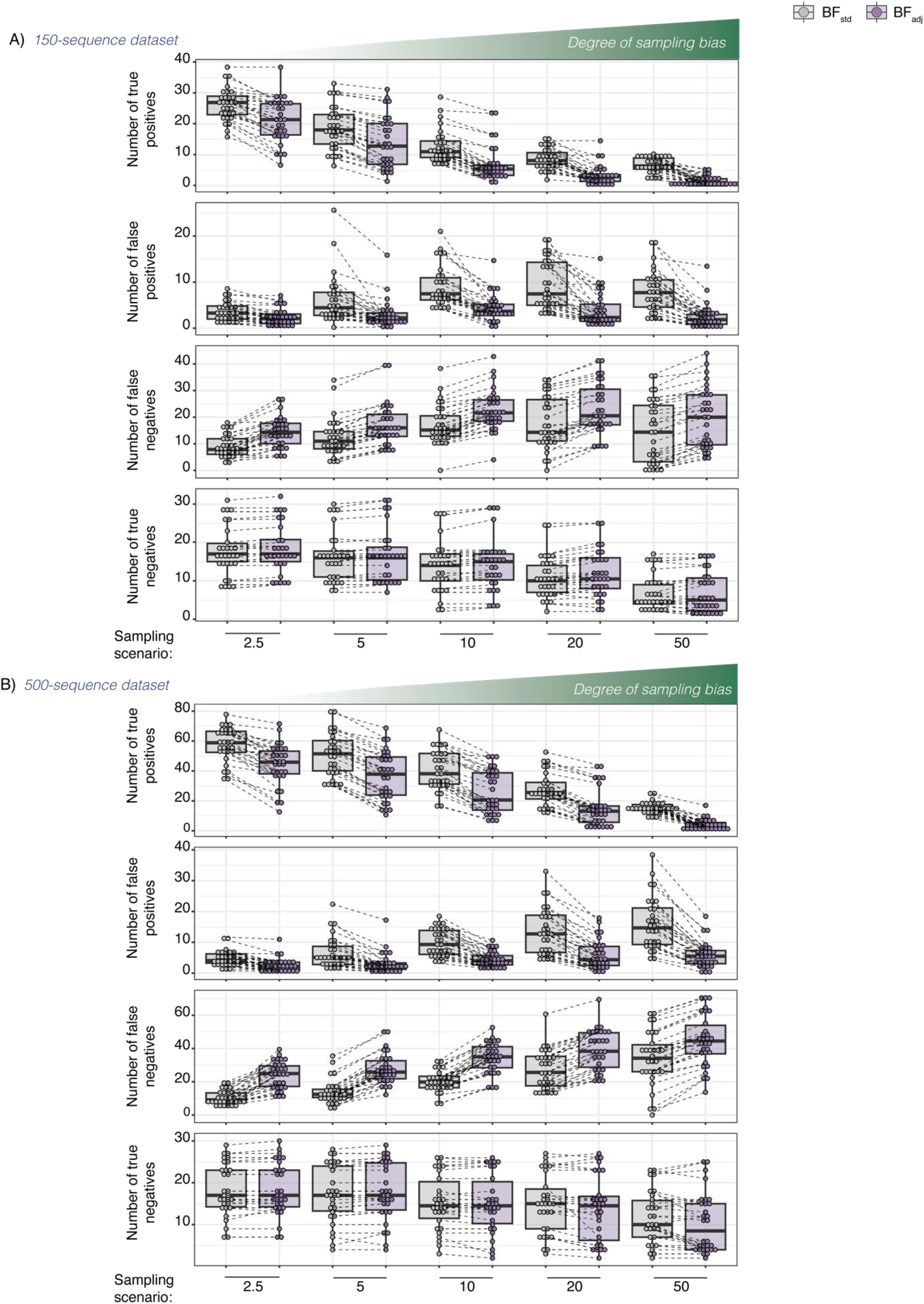
Impact of the **BF**_**adj**_ on the number of true positive (TP), false positive (FP), false negative (FN) and true negative (TN) inferred transition events under increasing levels of sampling bias using a cut-off value of 3. Panel A and B show box plots illustrating the distribution of the number of TPs, FPs, FNs and TNs inferred transition events across different sampling scenarios for 30 simulations, for the 150-sequence and 500-sequence datasets, respectively.

**Figure S3.**
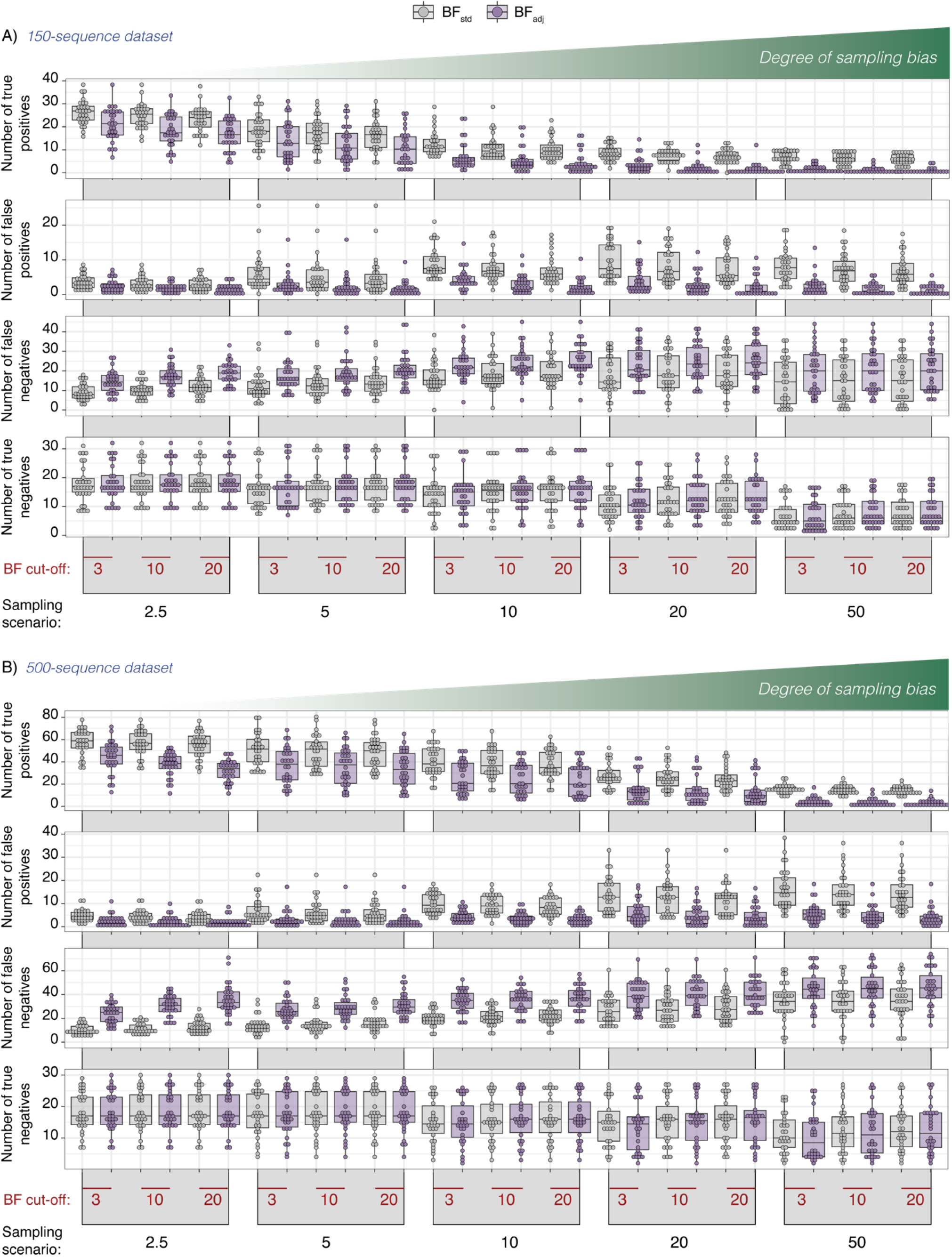
Impact of the **BF**_**adj**_ on the number of true positive (TP), false positive (FP), false negative (FN) and true negative (TN) inferred transition events under increasing levels of sampling bias and different BF cut-off values. Panels A and B show results for the 150-sequence and 500-sequence datasets, respectively. In each panel, the first row reports the TPs, the second row the FPs, the third row the FNs, and the fourth row the TNs. The x-axis has two levels: the first indicates different BF_std_ and BF_adj_ cut-off values (in red), while the second represents the different sampling scenarios (increasing sampling weights).

